# Comparative somatic genomics reveals divergent development of cell lineages across scleractinian corals

**DOI:** 10.64898/2026.05.19.726040

**Authors:** Trinity Conn, Thorsten B.H. Reusch, Benjamin Werner, Iliana B. Baums

## Abstract

Somatic mutations may drive adaptation and aging across diverse life forms, yet their role remains poorly understood in many early-branching animals. Here, we compare somatic mutation accumulation in the robust coral *Orbicella faveolata* with previous findings in the complex coral *Acropora palmata*. Whole-genome sequencing revealed high fixation of somatic genetic variants in *O. faveolata*, particularly in older, interior regions of colonies—contrasting with *A. palmata*. These patterns suggest distinct cell population dynamics between clades, indicating a segregated, mammal-like germline in *O. faveolata*, whereas such a germline remains undetected in *A. palmata*. This underscores the diversity of somatic evolutionary mechanisms across scleractinian corals.

## Background

Somatic evolution, where somatic mutations experience genetic drift and selection within an individual occurs across the tree of life, but its evolutionary implications have been studied primarily in vertebrates, largely in the context of aging and disease (1). In modular organisms such as plants, fungi, and cnidarians, Somatic Genetic Variants (SoGV) segregate within an individual through somatic genetic drift and selection because their simple, iterative body plans allow somatic mutations to segregate among ramets modules without harming the whole genet (2).

The fate of somatic mutations depends on their rate of occurrence, spatial organization of cell lineages, and selective pressure. In plants, a small number of meristem cells founding new shoots increases fixation probability (3). In corals, however, the number of stem cells contributing to new polyps and the degree of interconnection among polyps remain uncertain, limiting predictions about long-term somatic mutation dynamics.

Reef-building corals (Scleractinia) diverged into the major clades Complexa and Robusta approximately 415 million years ago (4–6). Although both groups include modular, colonial species (7,8), there are substantial functional and morphological differences which likely extend to somatic evolutionary processes. In the complex coral *Acropora palmata*, somatic mutations accumulate linearly with age but rarely reach fixation and can be transmitted to sexual offspring (9,10). In the robust coral *Orbicella faveolata*, limited data identified fixed somatic variants that did not appear to enter the germline (11). Here we tested the hypothesis that the rate of somatic fixation differs between the Robusta and Complexa clades which would point to variability in the underlying developmental architecture in these early branching metazoans

## Results and Discussion

*Orbicella faveolata* is a mounding coral, up to ∼3 m^2^ in size (12). Linear extension rates range from 0.6-0.8 cm per year with the polyps on the upper half of the colony contributing most to this growth (13-15). Here, we report the substitution rate, distribution of allele frequencies, and spatial segregation of somatic mutations within 3 colonies of *Orbicella faveolata*, each sampled along an upper and a lower transect (n = 4 samples / colony) and sequenced to an average read depth of 70x (Supplementary Table S1).

The mean substitution rate across colonies was 1.61 × 10^−4^ module^−1^ bp^−1^ approximately twice that reported for *A. palmata* (9) (Supplementary Table S2). Mutation load varied significantly among colonies (ANOVA, p < 0.05) and was consistently higher along upper transects, indicating greater accumulation in newly growing tissues. However, the proportion of fixed mutations was higher in lower transects (F = 42.31, p = 3.7 × 10^−9^), where older polyps are located. Interior modules had roughly ten-fold higher fixation than polyps at growth margins (Figure 1c.). While the proportion of fixation and substitution burden differed between colonies, likely due to difference in age (16), transect location explained significantly more variation in fixation than colony identity (two-way ANOVA, p<0.05). These results suggest that fixation is driven more by local cell-population dynamics than by overall colony age.

**Figure 1:**
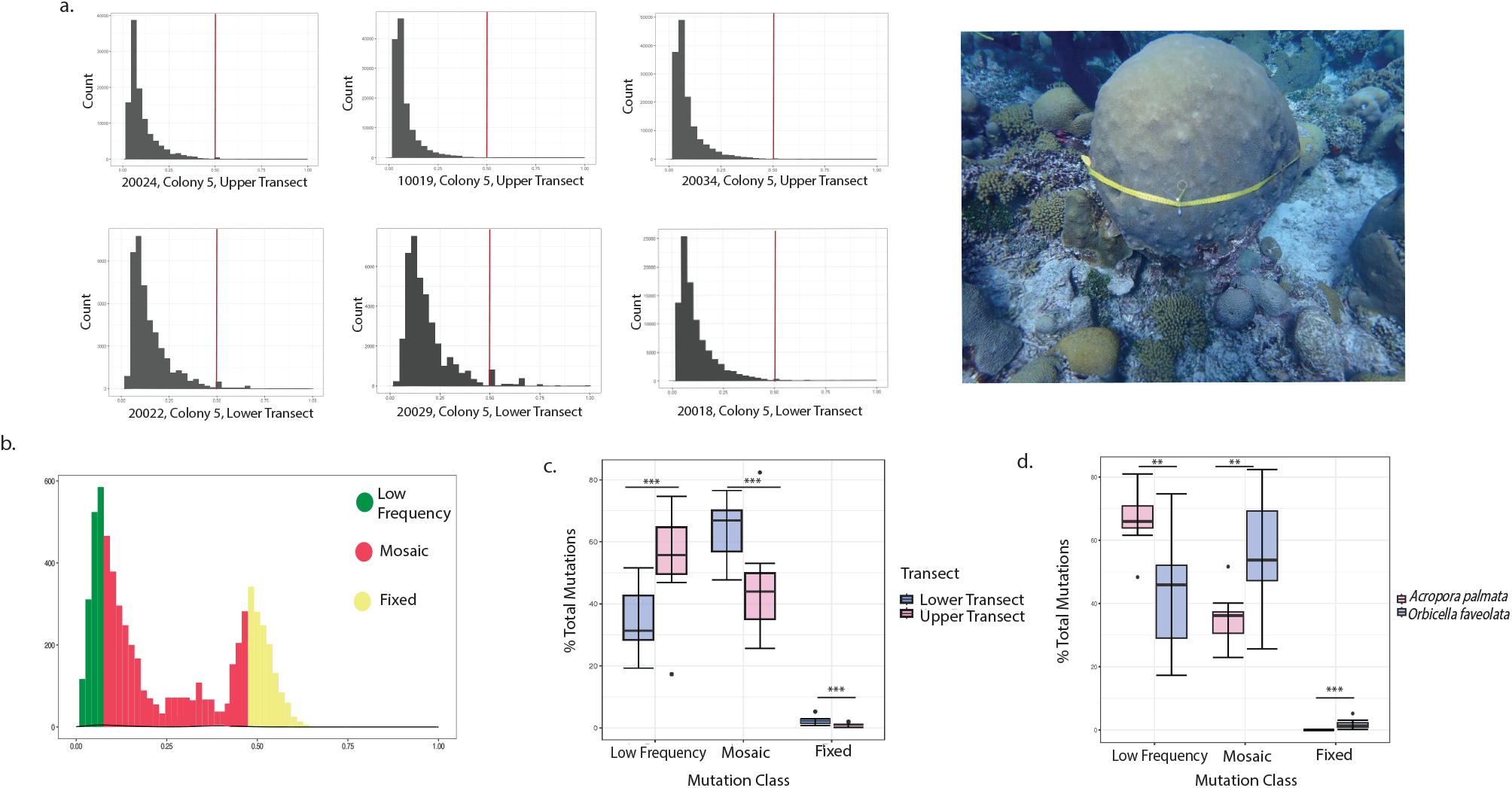
Patterns of Somatic Mutation Accumulation and Allele Frequency Within a Genet of *Orbicella faveolata*. (a) Allele frequency distributions of modules across colony 5 sampled in this study. Three samples were taken along the upper transect, and three across the lower transect. Image pictured is colony 5. Lower transect pictured. Red line in allele frequency distributions represents VAF=0.5, marking above where mutations were considered fixed.(b) Distribution of somatic mutations under a neutral model are categorized into three classes in this study. (c) Distribution of the three classes of mutations as the total proportion of mutations within a module. Mutations were classified as low frequency mutations (VAF= 0-0.1) mosaic mutations (VAF = 0.1-0.49) or fixed (VAF > 0.5). The proportion of mutations that fell into all three classes differed significantly (1-way ANOVA, p<0.05). The central line within each box represents the median of the percent of mutations, while end of the lines represent the range of the percent of total mutations. Lines with (***) designation indicate the significance between the lower and upper transect within each class of somatic mutation. (d) Distribution of the three classes of mutations as the total proportion of mutations within a module in *Orbicella faveolata* (in blue) and *Acropora palmata* (in pink). The proportion of mutations that fell into all three classes differed significantly between species (1-way ANOVA, p<0.05). The central line within each box represents the median of the percent of mutations, while end of the lines represent the range of the percent of total mutations. Lines with (***) designation indicate the significance between the species within each class of somatic mutation.

**Figure 2:**
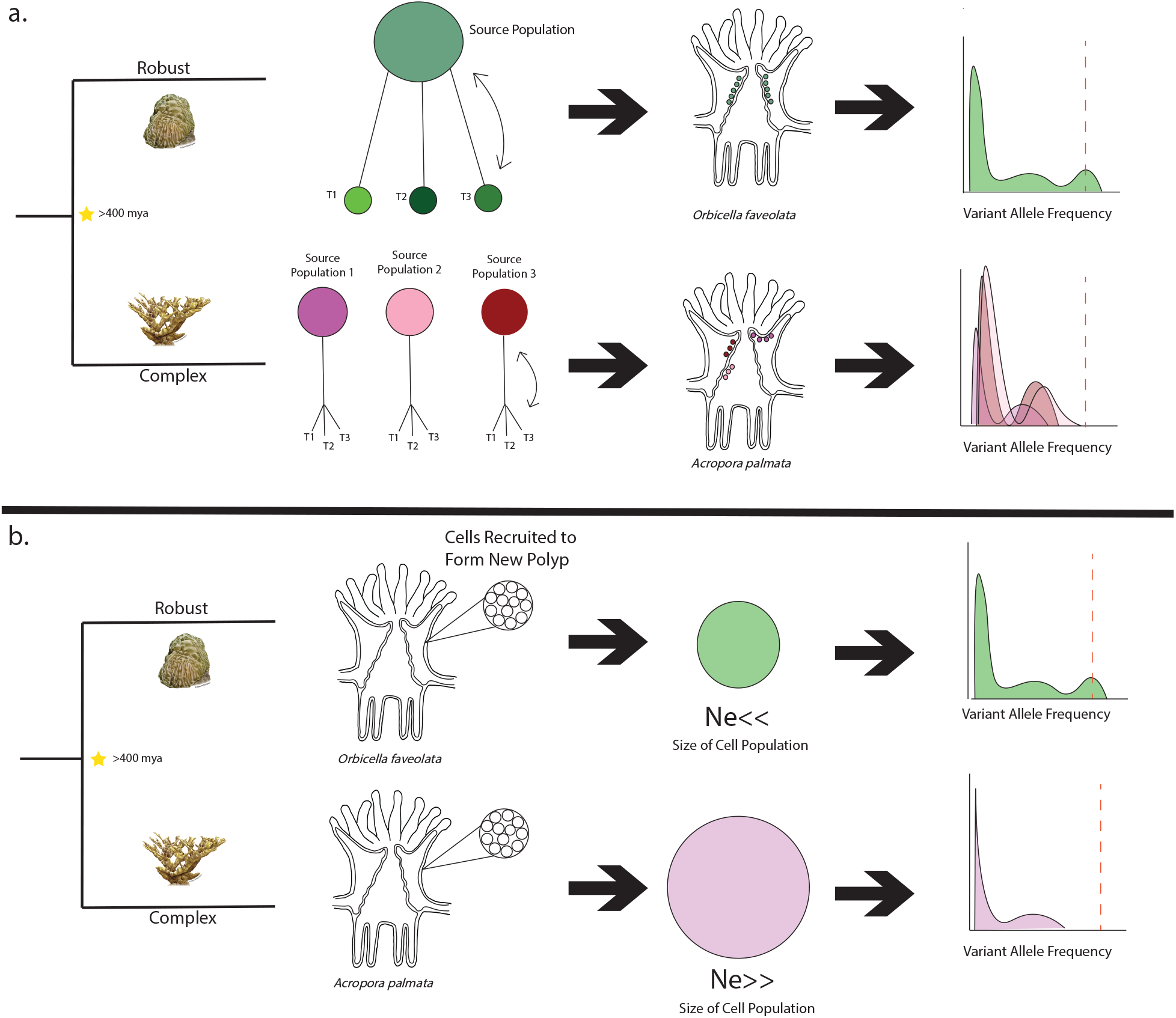
Hypothetical population dynamics to explain differences in allele frequency distributions and germline segregation in *Orbicella faveolata* and *Acropora palmata*. (a) Lack of fixation of somatic mutations in A. palmata could be explained by the presence of several independent stem cell lineages that each contribute cells to a new module. In any given sample, the observed allele frequency spectra would then be composites of several lineages. (b) Alternative hypothesis for variation is the size of the population recruited to produce a new polyp. Size of the stem cell pool size (here pictured as Ne) is a large contributor to the likelihood of fixation within an asexual population. If the stem cell pool is large, genetic drift will take longer to act. If A. palmata has a large stem cell pool, it could explain the lack of fixation observed. In contrast, *O*.*faveolata* may have a small effective population size of the cells contributing to the next polyp explaining the observed variant allele frequencies of mutations in this species. Sketches of *A*.*palmata* and *O*.*faveolata* are from NOAA.

These results contrast sharply with *A. palmata*, where somatic mutations remain in a mosaic state even in colonies estimated to be more than 100 years old (9). *O*.*faveolata* exhibited significantly higher proportions of fixed and mosaic SoGVs, whereas *A*.*palmata* contained proportionally more low frequency mutations (ANOVA, p<0.01. Theoretical models predict that fixation probability depends on both mutation rate and the number of foudning stem cells (3), consistent with these contrasting outcomes. Mosaicism in *A. palmata* may reflect a large, interconnected stem-cell pool which maintains mutations in a mosaic state across many modules. In contrast, extensive fixation in *O. faveolata* suggests either a smaller stem-cell pool, fewer founding cells per polyp that allow new modules to fix mutations by chance, higher mutation rates, or differences in genet ages.

While direct mutation rate estimates from cell cultures for *O. faveolata* are lacking, the elevated substitution burden observed in this study could reflect faster mutation accumulation. The evolutionary divergence between robust and complex corals exceeds that among mammalian orders known to differ in mutation rate (1, 17), suggesting that mutational processes themselves may have diverged across coral lineages.

Both *A. palmata* and *O*.*faveolata* depend on clonal reproduction with many genets persisting for centuries, fragmenting into ramets that continue to accumulate mutations independently, (9, 18-20) but no quantitative data exists on the relative distribution of genet ages between the species. Regardless of potential genet age difference between the sampled colonies of *A. palmata* and *O. faveolata*, the striking within-colony gradients in terms of fixed Somatic Genetic Variation (SoGV) in *Orbicella* suggest demographic differentiation among polyps in the latter that is not evident in *A. palmata*. Polyps at the center of an *O. faveolata* colony divide slowly and are effectively older, experiencing stronger drift and higher fixation, while polyps along the growth axes divide rapidly, maintaining larger effective populations that resist fixation The result is a spatially structured mosaic of evolutionary histories within a single *O. faveolata* colony (15). Mutations that arise in older polyps are more likely to become fixed locally, producing persistent sub clonal genetic structure that could influence physiological traits or local fitness within the colony.

The differences between *O. faveolata* and *A. palmata* highlight the importance of broad taxonomic sampling in studies of somatic evolution. Corals encompass exceptional diversity in morphology and life history, providing a natural experiment for understanding how multicellularity, clonality, and longevity shape evolutionary trajectories. Small differences in developmental organization, such as stem-cell bottleneck size, may alter the evolutionary role of somatic mutations.

## Conclusions

This study expands the genomic perspective on somatic evolution in basal metazoans. *O. faveolata* exhibits extensive within-colony fixation correlated to polyp age whereas *A. palmata* maintains long-term mosaicism. These contrasting patterns underscore the role of cellular population dynamics in determining how somatic variation contributes to organismal and evolutionary processes. Future studies incorporating additional species across clades alongside genets of known age will clarify how stem-cell demography and growth form interact to govern the evolutionary impact of somatic mutations in reef-building corals.

## Methods

### Study Design & Data Generation

The goal of this study was to quantify somatic mutation accumulation and fixation across spatially distinct modules within *O. faveolata* colonies. The study was conducted in May 2022 at the Water Factory reef site in Curaçao (12.10853° N, −68.95388° W). Three large colonies were sampled using pressurized drills. Each colony was divided into upper (actively growing) and lower (older interior) transects, with four tissue samples collected per transect. Samples were preserved in 95% ethanol and stored at −20 °C.

DNA was extracted using the Qiagen DNeasy Kit with modifications (Kitchen et al., 2020). DNA quality was verified by Nanodrop and agarose gel electrophoresis. TruSeq PCR-Free libraries (Illumina) were sequenced on a NovaSeq 6000 platform with 150 bp paired-end reads and a target coverage of 60–80× per sample

### Data Processing, Identification and Quantification of Somatic Burdan

Somatic mutations were identified using Mutect2 from the Genome Analysis Tool Kit (GATK) (21).Somatic mutations were identified as pairwise comparisons across all samples within a genet using a pairwise tumor-normal method.

Substitution rate was calculated per module as described in Conn *et al*., 2025. Within-colony variation of *O*.*faveolata* was quantified by comparing mean substitution rates between lower and upper transects.

Mutations were considered fixed in a module if the variant allele frequency of the mutation was ≥ 0.5. True fixation may be present as a gaussian curve around 0.5 due to sequencing error or contamination. Therefore, mutations that are classified as mosaic mutation may be fixed mutations; however, for the purposes of this study we used a conservative approach and only considered mutations that had at least a VAF of 0.5 as fixed. Mutations were classified as either low-frequency (VAF<0.1) mosaic (0.1 < VAF < 0.49) or fixed (0.5) (Figure 1b). The proportion of each mutation class was calculated as a fraction of the total unique mutation burden per sample. An ANOVA was used to test whether the proportion of mutations in each class was associated with the variability of testing individual genets (Colony) or their location within the genet (Transect). All statistical analyses were performed in R v.4.5.0.

## Abbreviations

SoGV: Somatic genetic variation
VAF: Variant Allele Frequency
bp: base pair
ANOVA: Analysis of Variance
GATK: Genome Analysis Toolkit

## Declarations

### Ethics approval and consent to participate

Not applicable

### Consent for Publication

Not applicable

## Acknowledgments

We thank Dr. Nicolas Locatelli for aid in fieldwork and sample collection and Zoe Dellaert for aid in fieldwork, sample collection, and input on figure design.

## Data availability

Scripts and associated data for this manuscript are available on Github https://github.com/TheBaumsLab/Orbicella-Somatic-Mutations). Raw sequence data is available on NCBI under BioProject PRJNA1243916.

## Permits

The research complies with applicable laws on sampling from natural populations and animal experimentation (including the ARRIVE guidelines). All samples were collected in Curaçao under permits provided to CARMABI by the Curaçaoan Government and were imported to the United States under CITES permit 1616US784243/67 where the samples were analyzed. No permitting requirements currently exist to govern access to corals under the Convention of Biological Diversity and its Nagoya Protocol or any other potentially applicable access and benefit sharing laws and regulations of Curacao.

## Funding

This work was funded from grant RGP0042/2020 awarded by the Human Frontiers Science Program to Drs. Iliana Baums, Thorsten Reusch and Benjamin Werner. Dr. Trinity Conn was supported by the Science Achievement Graduate Fellowship, the Human Frontiers Science Program, and through teaching assistantships at Pennsylvania State University during the production of this work.

## Author contributions

TLC conducted field work, DNA extraction and sample processing, bioinformatics and data analysis. IBB conducted field work. IBB, TR and BW acquired funding. TLC and IBB conceived the study. TLC and IBB wrote the manuscript. TLC, IBB, TR and BW edited the paper.

## Notes

### Competing Interest Statement

The authors have declared no competing interest.

## References

1. Cagan A, Baez-Ortega A, Brzozowska N, Abascal F, Coorens THH, Sanders MA, et al. Somatic mutation rates scale with lifespan across mammals. Nature. 2022;604(7906):517–24.

2. Reusch TBH, Baums IB, Werner B. Evolution via somatic genetic variation in modular species. Trends in Ecology & Evolution. 2021;36(12):1083–92.

3. Yu L, Renton J, Burian A, Khachaturyan M, Kotta J, Stachowicz JJ, et al. Precise age estimation in clonal species using a somatic genetic clock. bioRxiv. 2023:2023.11.07.566010.

4. Romano SL, Palumbi SR. Evolution of scleractinian corals inferred from molecular systematics. Science. 1996;271(5249):640.

5. Budd AF, Romano SL, Smith ND, Barbeitos MS. Rethinking the Phylogeny of Scleractinian Corals: A Review of Morphological and Molecular Data. Integrative and Comparative Biology. 2010;50(3):411–27.

6. Kerr AM. Molecular and morphological supertree of stony corals (Anthozoa: Scleractinia) using matrix representation parsimony. Biological Reviews. 2005;80(4):543–58.

7. Baums IB. A restoration genetics guide for coral reef conservation. Molecular Ecology. 2008;17(12):2796–811.

8. Lord KS, Lesneski KC, Buston PM, Davies SW, D’Aloia CC, Finnerty JR. Rampant asexual reproduction and limited dispersal in a mangrove population of the coral <i>Porites divaricata</i>. PROCEEDINGS OF THE ROYAL SOCIETY B-BIOLOGICAL SCIENCES. 2023;290(2002).

9. Conn T, Renton J, Chamberland VF, Dellaert Z, Reusch TBH, Werner B, et al. Mosaic accumulation of somatic genetic variation and estimates of age in the long-lived reef-building coral <em>Acropora palmata</em&gt. bioRxiv. 2025:2025.04.11.641527.

10. Vasquez Kuntz KL, Kitchen SA, Conn TL, Vohsen SA, Chan AN, Vermeij MJA, et al. Inheritance of somatic mutations by animal offspring. Science Advances. 2022;8(35):eabn0707.

11. Barfield S, Aglyamova GV, Matz MV. Evolutionary origins of germline segregation in Metazoa: evidence for a germ stem cell lineage in the coral Orbicella faveolata (Cnidaria, Anthozoa). Proceedings of the Royal Society B: Biological Sciences. 2016;283(1822):20152128.

12. E.H. Meesters, M. Hilterman, E. Kardinaal, M. Keetman, deVries, M., & R.P.M. Bak.(2001). Colony size-frequency distributions of scleractinian coral populations: spatial and interspecific variation. Marine Ecology Progress Series, 209, 43–54.

13. Sánchez-Pelcastre DW, Tortolero-Langarica JJA, Alvarez-Filip L, Cruz-Ortega I, Carricart-Ganivet JP. Sclerochronological characteristics of Orbicella faveolata in Cayo Arenas, a remote coral reef from the Gulf of Mexico. PLoS One. 2023;18(11):e0293802.

14. Ochoa-Serena A, Tortolero-Langarica JJA, Rodríguez-Troncoso AP, Carricart-Ganivet JP. Coral growth rates of the massive coral Orbicella faveolata in Puerto Morelos National Reef Park, Mexico. Regional Studies in Marine Science. 2025;89:104379.

15. Medellín-Maldonado F, López-Pérez A, Ruiz-Huerta L, Carricart-Ganivet JP. Understanding corallite demography to comprehend potential bias in sclerochronology: Analysis of coral modular growth by micro-computed tomography. Limnology and Oceanography. 2022;67(12):2665–76.

16. Yu L, Renton J, Burian A, Khachaturyan M, Bayer T, Kotta J, et al. A somatic genetic clock for clonal species. Nature Ecology & Evolution. 2024;8(7):1327–36.

17. Nei M, Xu P, Glazko G. Estimation of divergence times from multiprotein sequences for a few mammalian species and several distantly related organisms. Proceedings of the National Academy of Sciences. 2001;98(5):2497–502.

18. Devlin-Durante MK, Miller MW, Caribbean Acropora Research G, Precht WF, Baums IB. How old are you? Genet age estimates in a clonal animal. Molecular Ecology. 2016;25(22):5628–46.

19. Olsen KC, Moscoso JA, Levitan DR. Somatic Mutation Is a Function of Clone Size and Depth in Orbicella Reef-Building Corals. The Biological Bulletin. 2018;236(1):1–12.

20. Foster NL, Baums IB, Sanchez JA, Paris CB, Chollett I, Agudelo CL, et al. Hurricane-Driven Patterns of Clonality in an Ecosystem Engineer: The Caribbean Coral Montastraea annularis. PLoS One. 2013;8(1):e53283.

21. Van der Auwera GA, Carneiro MO, Hartl C, Poplin R, del Angel G, Levy-Moonshine A, et al. From FastQ Data to High-Confidence Variant Calls: The Genome Analysis Toolkit Best Practices Pipeline. Current Protocols in Bioinformatics. 2013;43(1):11.0.1-.0.33.

